# Virulence Network of Interacting Influenza-Host Protein Domains

**DOI:** 10.1101/2022.10.11.511722

**Authors:** Teng Ann Ng, Shamima Rashid, Chee Keong Kwoh

## Abstract

There exist several databases that provide virus-host protein interactions. While most provide curated records of interacting virus-host protein pairs, information on the strain-specific virulence factors or protein domains involved, is lacking. Some databases offer incomplete coverage of Influenza strains because of the need to sift through vast amounts of literature (including those of major viruses including HIV and Dengue, besides others). None have offered complete, strain specific protein-protein interaction records for the Influenza A group of viruses.

In this paper, we present a comprehensive network of predicted domain-domain interaction(s) (DDI) between Influenza A virus (IAV) and mouse host proteins, that will allow the systematic study of disease factors by taking the virulence information (lethal dose) into account. From a previously published dataset of lethal dose studies of IAV infection in mice, we constructed an interacting domain network of mouse and viral protein domains as nodes with weighted edges. The edges were scored with the Domain Interaction Statistical Potential (DISPOT) to indicate putative DDI.

The virulence network can be easily navigated via a web browser, with the associated virulence information (LD_50_ values) prominently displayed. The network will aid Influenza A disease modeling by providing strain-specific virulence levels with interacting protein domains. It can possibly contribute to computational methods for uncovering Influenza infection mechanisms mediated through protein domain interactions between viral and host proteins.

## 1 INTRODUCTION

Influenza A virus (IAV) is a single stranded, positive ribonucleic acid (RNA) virus that is a respiratory pathogen across many species such as humans, swine, and wild waterfowl. It consists of eight genomic segments which encode at least 11 proteins. The structure and organization of the virus particle is shown in Figure 1, which is reproduced here from the work of Jung and Lee [17].

**Figure 1.**
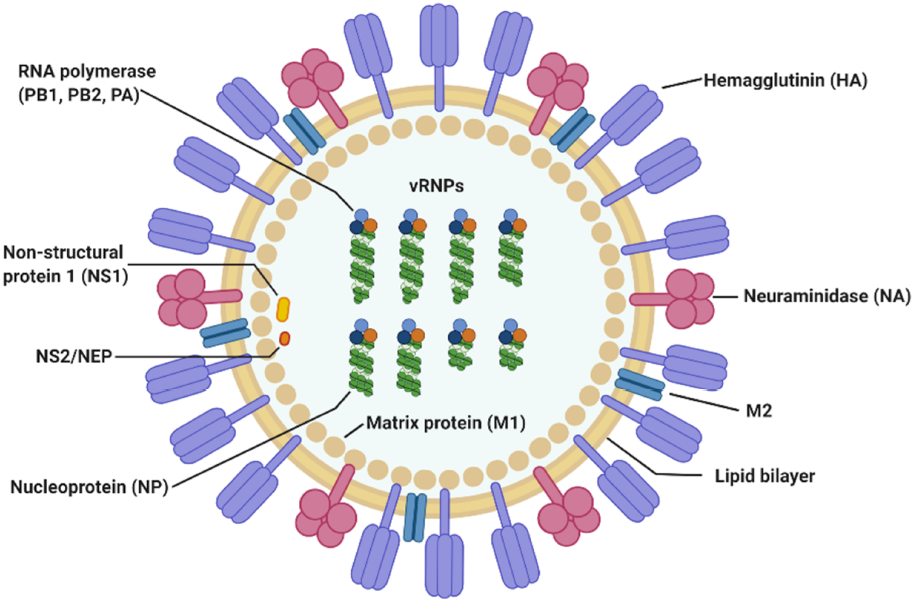
IAV particle and its fully assembled constituent proteins. Genomic RNA segments are shown in green, wrapped around nucleoproteins. The hetero-trimeric RNA-dependent RNA polymerase complex comprising of PB1, PB2 and PA is shown in orange, light and dark blue circles. This figure is by Jung and Lee [17].

Hemagglutinin (HA) and neuraminidase (NA) on the viral particle surface, are the proteins responsible for mediating entry into and cleavage from the host cell, respectively. Matrix protein 1 (M1) is a component of the viral envelop while matrix protein 2 (M2) is found below the lipid bilayer of the viral membrane, strengthening it. Together with the nucleoprotein (NP), they form the ribonucleoprotein complex (indicated as vRNP in Figure 1). The final three proteins are the polymerase basic 1 frame 2 (PB1-F2), and non-structural proteins 1 and 2 (NS1 and NS2), respectively.

IAV can be highly pathogenic in humans and several highly virulent strains have already caused millions of deaths worldwide in multiple pandemic events. Estimated death tolls for the 1918 (H1N1), 1957 (H2N2) and 1968 (H3N2) pandemics are 50 million [16], 1.1 million [37] and 1-4 million [29], respectively. Further, IAV triggers various respiratory illnesses seasonally, making it endemic in human populations. The yearly number of deaths due to influenza associated respiratory illness from seasonal influenza has been estimated to vary between nearly 300,000 to 646,000 [14]. Since it is endemic in populations, hosts harboring seasonal influenza strains can act as a reservoir for reassortment events, leading to cross-infection with other circulating pathogens such as SARS-CoV-2 to form potentially harmful recombinant strains [35]. These attributes highlight the complexity of disease factors of respiratory pathogens and indicate the need of wide-scale influenza studies. They also make the continual monitoring of public health outcomes necessary.

Infectious studies of mouse models can help to elucidate host factors responsible for virulence, since they are cost effective, reproducible and allow mechanistic analyses that may not be directly conducted on humans due to ethical reasons [30]. One way to measure the pathogenicity of IAV is by obtaining the lethal dose at which 50% of the inoculated animal test population is infected or perishes (abbreviated here as LD_50_) [11]. By comparing outcomes of Influenza infections in different strains of mice, differences due to allelic variations in mice strains could be possibly be established [23].

Data records of mouse model infectious studies had been previously collected in an earlier work by F.X. Ivan and C.K. Kwoh [15]. Their study highlighted the role of protein sites of PB2 in Influenza virulence by a systematic meta-analysis using rule-based models to predict the virulence. Therefore, a link between macroscopic virulence labels (such as LD_50_ categories) and protein-protein interactions could prove beneficial in understanding the factors contributing to IAV virulence. Domain-domain interaction(s) DDI can be particularly useful because a protein domain is often a discrete functional unit that is modular, and protein-protein interactions rely on combinations of DDI [13, 2]. Hence here, the network was constructed with domains representing nodes. While ’domain-domain’ interactions are by definition a subset of ’protein-protein’ interactions, here the quoted terms are used interchangeably, unless specified otherwise.

To systematically identify potential interactions between IAV and mouse host-proteins, a protein network consisting of putative DDI between IAV and mouse proteins, scored by the Domain Interaction Statistical Potential (DISPOT) [24] was developed in this work.

The protein network is presented in a clear graphical user interface (GUI) that easily shows the LD_50_ values and interacting protein domains from the C57BL/6J mouse strain as identified by DISPOT. The virulence network of interacting protein domains will assist studies of IAV disease modeling by providing data of putative interacting protein domains that are associated with their LD_50_ values.

The rest of this manuscript is organized as follows. Section 2 details the contents and design and describes the data presented in this database and the data curation procedure. Section 3 details the web server implementation, describing the tools used, graphical user interface layout and functionality. Section 4 details discussion of this data. Section 5 outlines the proposed future work. Section 6 summarizes and concludes this paper.

## 2 CONTENTS AND DESIGN

Figure 2 outlines the steps taken to the implement IAV-Host protein–protein interaction (PPI) database. The IAV-Host PPI web server can be accessed at: https://iav-ppi.herokuapp.com/home.

**Figure 2.**
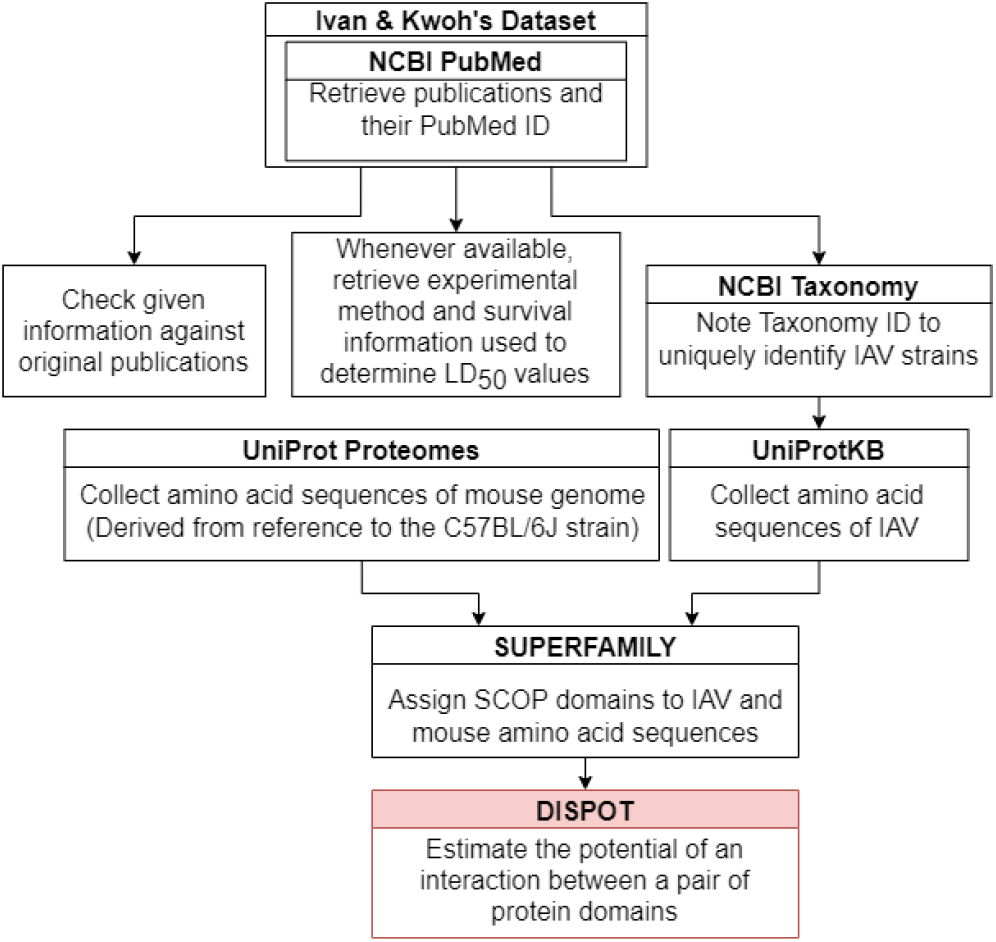
Implementation procedure.

The IAV-Host PPI database provides information of host-pathogen interactions between 109 unique IAV strains across 14 IAV subtypes and 2419 unique mouse proteins. Five out of the eight RNA segments of IAV genome, namely PB1, HA, NA, M1 and NS1 were found to contain the interacting pathogen protein domains. In summary, 31 pairs of DDIs were found between seven IAV protein domains and 29 mouse protein domains.

This work builds on initial data records of mouse model infectious studies collected in the previous work by F.X. Ivan and C.K. Kwoh [15]. Their same process of assigning virulence levels was followed here. LD_50_ value was the, key information needed to identify the virulence class of a specific IAV strain. Virulence was classified as two-class (avirulent/virulent) and three-class (low/intermediate/high) (*shown in Table 1).* Essentially, the total infection records classified as ‘virulent’ under the two-class problem is the sum of records classified as ‘intermediate’ and ‘high’ under the three-class problem. Likewise, the total infection records classified as ‘avirulent’ is equivalent to the number of records classified as : ‘low’ *(in Supplementary Figure S1).* For the three-class virulence classification, LD_50_ thresholds of 10^3.0^ and 10^6.0^ were applied, referencing thresholds that are used by World Health Organization (WHO), for classification of influenza virulence in mice (EID_50_ infection unit) [1]. LD_50_ infection units include Plaque-forming Unit (PFU), Focus-forming Unit (FFU), median (50%) Egg Infective Dose (EID50), 50% Tissue Culture Infectious Dose (TCID50) and 50% Cell Culture Infectious Dose (CCID50), where the equality across all units was assumed.

**Table 1.** Three-class virulence classification

| Low | Intermediate | High |
| --- | --- | --- |
| $LD_{50} > 10^{6.0}$ | $LD_{50} \leq 10^{6.0}$<br>and $LD_{50} > 10^{3.0}$ | $LD_{50} \leq 10^{3.0}$ |

SUPERFAMILY 2.0 sequence search (https://supfam.org/sequence/search) [26] was used to map regions of an amino acid sequence to at least one Structural Classification of Proteins (SCOP) superfamily using the SUPERFAMILY hidden Markov models. SCOP is a representation of structure-based hierarchical classification of relationships between protein domains, with ‘family’ being the first level and ‘superfamily’ being the second level. Protein domains from the same SCOP family are strongly related and frequently share the same function [3].

Domain Interaction Statistical Potential (DISPOT) (http://dispot.korkinlab.org/home/pairs) [24] served as the web tool to determine presence of DDIs between pairs of IAV and mouse SCOP superfamily domains. DISPOT uses exclusively DDIs from DOMMINO [20], an in-depth database of structurally resolved macromolecular interactions, where data about DDIs is the amplest, as its source of data. For a given domain pair, DISPOT returns a statistical potential, denoted as the probability Pij. Statistical potentials take values across the entire scale of real numbers. Negative and positive values can be respectively interpreted as having more or less than average number of DDIs in the DOMMINO database. Neutral values are corresponding to the number of DDIs close to the average number. “No information” will be returned instead of a numeric value if the DOMMINO database does not have an entry for the particular domain pair.

The DISPOT calculation of statistical potential formula is given in the equation as follows [24]:

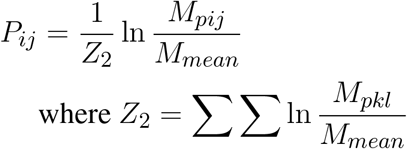

Z_2_ is the natural logarithm of observed frequencies of interactions between domains in the DOMMINO database. Mmean is the average number of interactions for a pair of domain families, calculated from the non-redundant DOMMINO dataset. Non-redundant refers to two corresponding pairs of domains that do not share 95% or more sequence identity [24].

### 2.1 Dataset

In this work, 57 journal publications were retrieved from National Centre for Biotechnology (NCBI) PubMed [22]. In all these publications, LD_50_ values were explicitly stated. 55 publications referenced supplementary information given in F.X. Ivan and C.K. Kwoh’s publication (“Additional file 5: Table S1”) [15], where LD_50_ values were stated as ”values given”. LD_50_ values reflected in their dataset were checked against the original publications and some missing records were added. Additionally, seven new records from two other papers [33] and [34] were documented.

#### 2.1.1 Data cleaning

The preliminary dataset presented in this work (https://github.com/tengann/IAV-Host-PPI-Database/blob/main/RawData_2022.xlsm) holds 488 infection records involving wild-type, laboratory, mouse-adapted, recombinant or mutant IAV strains. IAV genomes of wild-type strains are in their natural and non-mutated form while laboratory strains were prepared by means of reverse genetics. Mouse-adapted strains were derived from serial lung-to-lung passages of virus in mice. Genetic amino acid sequences of mutant virus were changed through point mutations via single amino acid substitutions. Recombinant strains were formed by the combination of protein segments from at least two different IAV strains.

The initial dataset was manually curated to only include records involving wild-type or laboratory IAV strains, thereby reducing the number of infections to 190 *(Supplementary Figure S2).* Subsequently, infection records comprising wildtype or laboratory strains where their Taxonomy identification (ID) number (otherwise known as accession number) [31] could not be found were dropped, further reducing the records to 166 *(Supplementary Figure S3).* In these cases, it was not possible to retrieve the complete protein sequences of IAV gene segments for SCOP domain assignment via SUPERFAMILY 2.0.

From the above process, the tally of infection records was reduced to 139 *(Supplementary Figure S3)*. Multiple records concerning the same combination of IAV strain and mouse genome were condensed into a single record, adopting the approach from F.X. Ivan and C.K. Kwoh’s publication [15]. Whenever possible, the majority class of the three-class virulence assignment scheme was selected. Otherwise, the class that is more or most virulent was considered. Next, if only the lower bound of the LD_50_ value was presented, the record with the highest lower bound was selected. For cases where the lowest exact or upper bound of LD_50_ value was provided, the record was selected. The final cleaned dataset contains 109 unique records and was used to derive the network of interacting protein domains.

### 2.2 Data Annotation

Firstly, to distinguish between all journal publications referenced, the NCBI PubMed (https://pubmed.ncbi.nlm.nih.gov/) [22] ID number uniquely assigned to each publication record was noted. Information collected in F.X. Ivan and C.K. Kwoh’s dataset consists of IAV strain, mouse strain, LD_50_ value and infection unit. Then, in this paper, to provide a deeper insight into how the LD_50_ value was determined in each separate experiment, additional evidence, namely, the experimental method, weight loss and/or survival remarks and LD_50_ calculation method were documented. Also, for each IAV strain, the Taxonomy ID number, a unique ten-digit code that designates classification and specialization was retrieved from NCBI Taxonomy database (https://www.ncbi.nlm.nih.gov/taxonomy) [31].

Amino acid sequences of both IAV and mouse proteins were retrieved from the UniProt (release 2021_03) protein knowledgebase (UniProtKB) (https://www.uniprot.org/) [5]. IAV protein sequences were retrieved using strain names and/or matching Taxonomy ID number where available. Mouse protein sequences were retrieved using the Proteome ID number, UP000000589. This reference proteome was derived from the genome sequence of mouse strain C57BL/6J, with Taxonomy ID number, 10090. For this work, among 55, 315 protein records that were available, 17, 120 Swiss-Prot gold star reviewed entries (https://www.uniprot.org/uniprotkb?query=UP000000589) were retrieved. Swiss-Prot reviewed refers to records with information fully and manually extracted from literature or curator-evaluated computational analysis [4]. It strives to provide high-quality annotations with a minimal level of redundancy and high level of integration with other databases.

As an average protein consists of two or more domains, domain start and end residue numbers, corresponding to regions of protein sequences matching each assigned SCOP domain (by SUPERFAMILY 2.0) were independently noted for every protein sequence retrieved from UniProtKB. Since each domain has its distinct structure and biological function, only a subset of domains constituting each protein are involved in the interaction between a pair of proteins. Thus, this enhances the complexity of host-pathogen protein interaction analysis [24].

Overall, IAV strains in the dataset were found to comprise proteins domains belonging to 13 SCOP superfamilies *(Table 2).* Then, these domains were paired up individually with 1102 unique domains found among the mouse protein sequences, forming a total of 14, 326 IAV-mouse protein domain pairs. Subsequently, domain pairs were fed as input into DISPOT, for calculation of statistical potential. Finally, seven domains in IAV proteins (indicated in red in *Table 2)* and 29 domains in mouse proteins were found to be involved in the host-pathogen PPI. Out of the 17, 120 mouse protein sequences retrieved, 2419 unique proteins were found to contain to at least one of the 29 interacting SCOP domains. In addition, the mouse protein localization in vital organs (lungs, brain, liver, kidney, spleen and heart) or blood was noted, whenever available. *(available in Supplementary Figure S4).*

**Table 2.** IAV Domains identified by SCOP Superfamily. Red indicates domains identified as interacting with mouse proteins, while - indicates no identified domains.

| IAV Segment | SCOP Superfamily name / Accession number |
| --- | --- |
| PB2 | PB2 C-terminal domain-like / 160453 |
| PB1 | DNA/RNA polymerases / 56672 |
| PB1-F2 | - |
| HA | Viral protein domain / 49818<br>Influenza hemagglutinin (stalk) / 58064 |
| NP | Flu NP-like / 161003 |
| NA | Sialidases / 50939 |
| M1 | Influenza virus matrix protein M1 / 48145<br>Alpha-catenin/vinculin-like / 47220<br>Methyl-accepting chemotaxis protein (MCP) signaling domain / 58104 |
| M2 | - |
| NS1 | NS1 effector domain-like / 143021<br>S15/NS1 RNA-binding domain / 47060 |
| NS2 | Nonstructural protein NS2, NEP, M1-binding domain / 101156<br>Spectrin repeat / 46966 |

## 3 WEB SERVER IMPLEMENTATION

IAV-Host PPI web server GUI has a comprehensible interface, made up of two pages, with various features, including browsing via subtype and strain to view information collected from literature searches, an interactive network graph with accompanying information on node and edge attributes as well as amino acid sequences extracted from UniProt. Figure 3 illustrates navigation and layout of the web server’s GUI.

**Figure 3.**
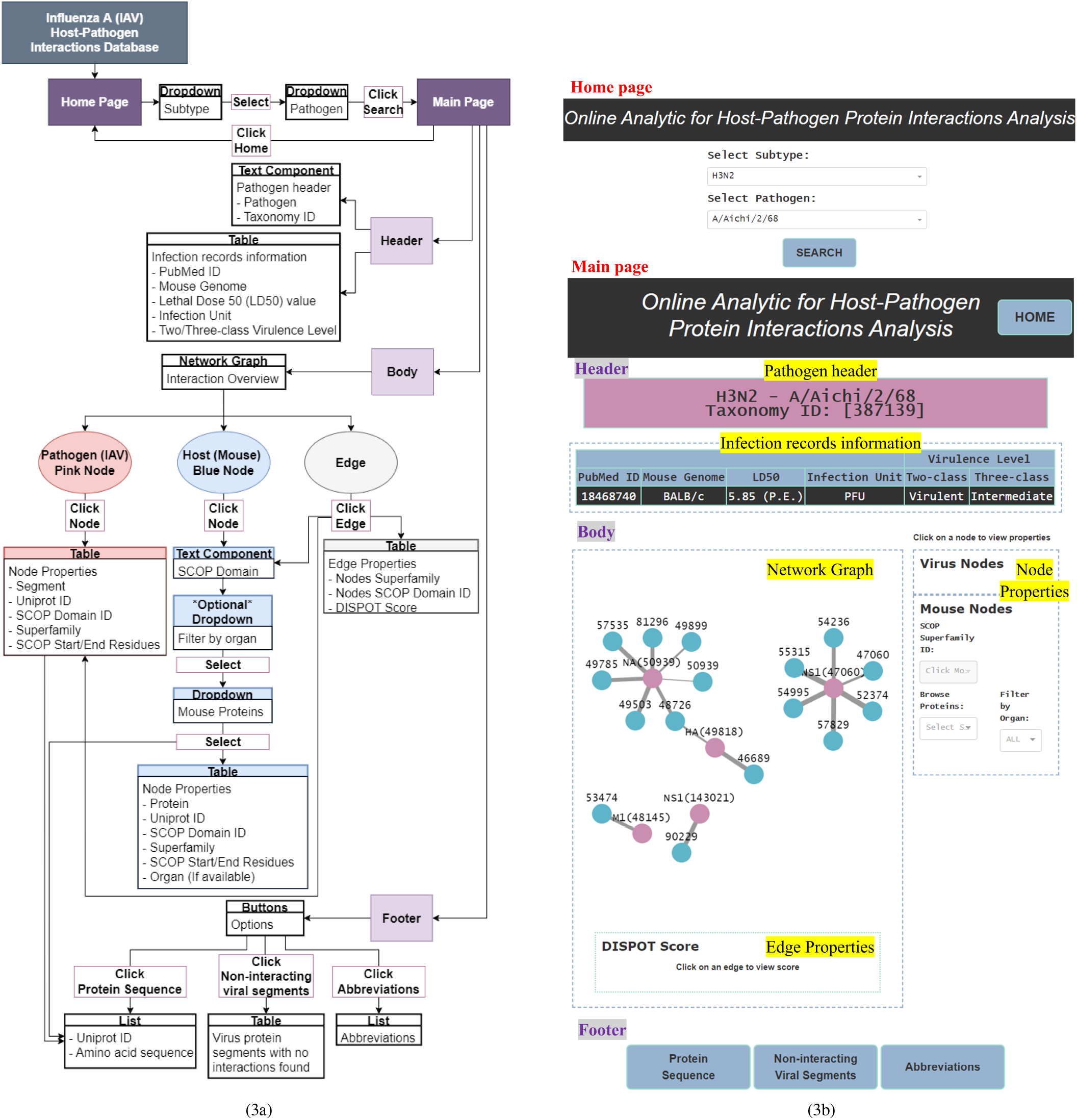
**(a)** GUI functionality navigation. **(b)** Screenshot of GUI layout.

### 3.1 Tools

Firstly, Microsoft Excel 2016 was used to store and organize data collected from literature. Secondly, the web interface was developed on the code editor Visual Studio Code V1.71.2, with Python V3.7.4 as the programming language. Python libraries used were Pandas V1.3.5, for transforming comma-separated values (csv) from Excel files to dataframes. Dash V2.3.1 was the framework for designing the application’s functionalities, described as follows: Dash bootstrap components V0.3.0 for building the application’s layout, graph visualization component Dash Cytoscape V0.3.0 for constructing the interactive network graph. Beautiful Soup V4.11.1 was the HTML parser used for pulling protein sequences from the UniProt database. Lastly, the cloud platform Heroku V7.62.0 was utilized to build and run the web interface entirely in the cloud.

### 3.2 Graphical User Interface (GUI)

#### 3.2.1 Home page

The home page features a dependent dropdown component to firstly allow the user to search for a specific IAV subtype and subsequently, browse and select the pathogens belonging to the selected subtype group.

#### 3.2.2 Main page

The main page features three main sections – header, body and footer (shown in Figure 3(b)).

##### 3.2.2.1 Header

The header section consists of two components – pathogen header text component and infection records information table.

###### Pathogen header

This text component displays the IAV subtype, pathogen name and Taxonomy ID, based on selections made by the user in the home page.

###### Infection records information

This section presents infection records information collected directly from literature and additional information collected from web tools, NCBI Taxonomy and UniProt databases in the form of a table. Information is filtered to present only those relevant to the user’s selections in the home page.

##### 3.2.2.2 Body

The body section consists of two panels, where the left panel displays the network graph and edge properties. The right panel is divided into two subsections and displays the virus and mouse node properties, respectively.

###### Network graph

The network graph was designed such that the user can clearly differentiate between IAV and mouse nodes by colors, where pink was assigned to IAV nodes and blue to mouse nodes. User can differentiate interaction statistical potentials by edge weights, where a thicker edge line represents a higher possibility of interaction. Also, the edge color will change to blue upon clicking, to highlight the edge selection.

###### Node and edge properties

Node properties include either the IAV protein segment or name of mouse protein, UniProt ID, SCOP superfamily ID, superfamily name and SCOP start/end residue(s). Edge properties comprise IAV and mouse SCOP superfamily ID and name with the matching DISPOT statistical potential score. All IAV node properties will be populated upon clicking of any pink IAV node while only the mouse SCOP superfamily ID field will be populated upon clicking of any blue mouse node in the network graph. Similarly, the former, together with its respective edge property will be presented upon click on any edge. To display all mouse node properties, a mouse protein first needs to be selected from the ”Browse Proteins” dropdown under the mouse node properties subsection.

##### 3.2.2.3 Footer

The footer provides the user with the following supplementary information – protein sequence, noninteracting viral segments and abbreviations.

###### Protein sequences

Web scraping was applied to extract amino acid sequences of IAV and mouse proteins from the UniProt database.

###### Non-interacting viral segments

Non-interacting IAV protein segments consist of the following cases: (1) The UniProt ID could not be found. Therefore, the protein sequence for input to SUPERFAMILY 2.0 is unknown and no domain information could be retrieved. In this case, the UniProt ID field was indicated with ”Not Found” and remaining information was labelled as “N/A”. (2) Some protein sequences retrieved were not mapped to any SCOP superfamily based on the SUPERFAMILY 2.0 database. As such, there was no protein domain information for input to DISPOT. (3) IAV domain information was available but the DOMMINO database did not have any entry between the IAV domain and all of 17, 120 retrieved mouse proteins, hence no interaction information was returned by DISPOT.

###### Abbreviations

This section conveys extra information to the user; specifically, the definitions and expansions of abbreviations used as well as notes targeted to help the user better comprehend the network graph.

## 4 DISCUSSION

This section lists domain pairs identified with high scoring interaction potentials. In section *4.1* the interacting domain pairs are verified against actual protein-protein interactions identified in biochemistry or proteomics literature. It is organized as a series of discussions on the functional role of interacting domains pairs, per paragraph.

### 4.1 Interacting protein domains

According to DISPOT scores obtained, the Top 10 domain pairs with strongest statistical interaction potentials are as listed in *Table 3.*

**Table 3.** Top 10 Interactions according to statistical potentials returned by DISPOT. A more negative DISPOT score indicates a higher possibility of interaction.

| IAV Segment | IAV SCOP domain | Mouse SCOP domain | DISPOT Score |
| --- | --- | --- | --- |
| NS1 | S15/NS1 RNA-binding | Nucleotidyl transferase | -4.301855801 |
|  |  | L30e-like |  |
| PB1 | DNA/RNA polymerases | DNA clamp |  |
|  |  | 5' to 3' exonuclease, C-terminal subdomain |  |
|  |  | PIN domain-like |  |
|  |  | N-acetylmuramoyl-L-alanine amidase-like |  |
| NS1 | S15/NS1 RNA-binding | Zn-binding ribosomal proteins | -3.608708621 |
|  |  | Ribosomal protein S6 |  |
| M1 | Alpha-catenin/vinculin-like | PH domain-like |  |
|  |  | I/LWEQ domain |  |

Of 109 unique IAV strains presented in this database, the PB1 gene segment of three strains *(Table 4)* were assigned the DNA/RNA polymerases domain. However, despite the strong interactions, it was not possible to ascertain if the presence of the DNA/RNA polymerases domain has an impact on the pathogenicity, due to variation in virulence levels across the different IAV strains.

**Table 4.** IAV strains with DNA/RNA polymerases domain assigned to PB1 gene segment.

| IAV Strain | Subtype | Three-class virulence level |
| --- | --- | --- |
| A/PuertoRico/8/1934 | H1N1 | High (BALB/c, DBA/2)<br>Intermediate (C57BL/6J) |
| rA/X-31 | H3N2 | Low (BALB/c) |
| A/HongKong/97/98 | H5N1 | Low (BALB/c) |

The M1 gene segment of eight IAV strains were assigned a Methyl-accepting chemotaxis protein (MCP) signaling domain instead of alpha-catenin/vinculin-like. No interaction was found between the MCP signaling domain with all domains present in mouse proteins. Comparing results from experiments carried out in H3N2 IAV strains *(Table 5 and 6),* especially on mice strains BALB/c and DBA/2, it is evident that presence of the alpha-catenin/vinculin-like domain is a virulence factor responsible for IAV infection. Alpha-catenins are members of the vinculin family of proteins. Vinculin is an actin-binding protein. Protease treatment revealed that actin present in the interior of influenza virions presumably participates in moving viral components to the assembly site and cytoskeletal reorganization that occurs during bud formation [27]. Actin is a family of globular multi-functional proteins that form microfilaments in the cytoskeleton. The host cytoskeletal network takes part in transport of viral components in the cell, predominantly during the stages of virus entry and exit [32].

**Table 5.**
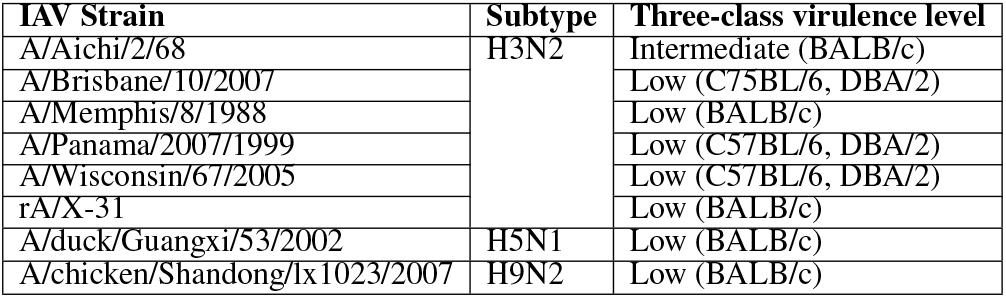
IAV strains with a Methyl-accepting chemotaxis protein (MCP) signaling domain assigned to M1 gene segment instead of alpha-catenin/vinculin-like.

**Table 6.**
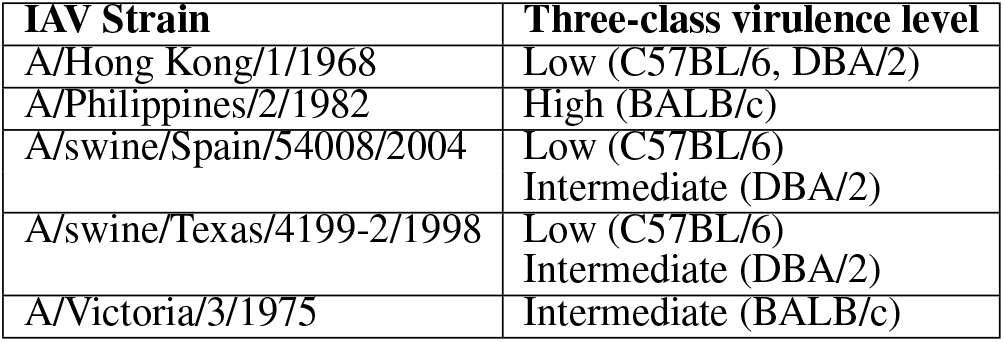
H3N2 strains with alpha-catenin/vinculin-like domain assigned to M1 gene segment.

Pleckstrin homology (PH) domain-like is a short peptide module often found in cytoskeletal proteins [41]. Although cytoskeletal elements are known to be associated with M1, the underlying mechanisms are not clear [43]. Ezrin, *[EZRI MOUSE (UniProt accession number: P26040)]* has a PH-like domain. Based on meta-analysis of IAV interactome studies on the M1 gene segment conducted by [7], Ezrin was discovered to be a common interactor and is a positive regulator of virus replication.

Information from UniProtKB indicates that majority of proteins that contain a nucleotidylyl transferase domain possess the tRNA ligase enzyme, otherwise known as aminoacyl-tRNA synthetase (ARSs). ARSs play a crucial role in protein synthesis by attaching amino acids to their cognate transfer RNAs (tRNAs) [25]. Specifically, Cysteine–tRNA ligase *[SYCC MOUSE (UniProt accession number: Q9ER72)],* an interactor of NS1, catalyzes the ATP-dependent ligation of cysteine to tRNA (Cys) and plays a role in translation [9]. Furthermore, ARSs plays a vital role in the development of immune cells because of their involvement in maturation, transcription, activation, and recruitment of immune cells. More significantly, ARSs regulate various biological processes and act as signaling molecules in infectious disease [25], which supports the high DISPOT score (−4.30) predicted for the S15/NS1 RNA-binding domain in NS1 segments.

Ubiquitin-40S ribosomal protein S27a, *[RS27A_ MOUSE (UniProt accession number: P62983)],* is a protein with the Zn-binding ribosomal protein domain. It is an NS1-interacting host protein node, classified as belonging to the apoptosis pathway [36]. Although not required for ribosome function, it plays an important role in the life cycle of IAV through regulating viral nucleic acid replication and gene transcription. When interrupted in host cells, the replication and infectivity of IAV is stopped [21].

CCCH zinc finger present in mouse proteins is the sole domain that interacts with NS1 effector domain-like instead of the S15/NS1 RNA-binding domain in the NS1 segment of IAV. This domain pair has a DISPOT statistical potential score of - 2.916 *(rounded to 3 d.p.).* Mouse protein, cleavage and polyadenylation specificity factor subunit 4 (CPSF4) *[CPSF4_MOUSE (UniProt accession number: Q8BQZ5)]* contains the CCCH zinc finger domain. The interaction between NS1 and CPSF4 controls the alternative splicing of tumor protein p53 (TP53) transcripts, and alters the expression of TP53 isoforms in parallel. As a result, cellular innate response, particularly via type I interferon secretion is regulated, leading to efficient viral replication [10].

The Immunoglobulin (Ig) domain, otherwise known as antibodies is the sole SCOP protein domain that interacts with three IAV domains, namely DNA/RNA polymerases, Viral protein domain and Sialidases in the PB1, HA and NA segments respectively. Immunoglobulin is the most abundant domain found among the 2419 unique mouse proteins containing interacting SCOP domains *(Supplementary Figure S4)*. During natural infection with IAVs, immune response against both HA and NA will be evoked [8]. IgM response is dominant in primary infection, while IgG response is dominant in secondary infection, for Ig secretion. IgA present in nasal secretions can neutralize HA and NA of IAVs [6].

In Tables 4, 5 and 6, BALB/c, C57BL/6J and DBA/2 denote the different mouse strains that were used in separate experiments.

### 4.2 Non-interacting protein domains

The non-interacting influenza hemagglutinin (stalk) SCOP domain present in the HA is an example that not all domains constituting a protein are involved in interaction between a pair of proteins. The stalk evolves slower than the receptor binding head and it is suggested that it has to remain structurally conserved owing to its role in membrane fusion [19]. Studies have also suggested that the stalk domain is not under immune pressure [38, 28]. Additionally, mutations in the stalk domain do not drastically impact virus binding or aid in avoiding neutralizing antibody responses from the host [19]. Therefore a potential ’true negative’ interaction is also identified in the protein domain network, in line with experimental findings.

### 4.3 Limitations

Generally, DISPOT works as a tool to streamline the PPI prediction problem through providing insight on the possibility of specific DDIs in a given physical PPI. However, it is not a classification method and statistical potentials returned are useful for ranking DDIs but do not directly translate to the probability score. DISPOT which solely uses information about interactions between protein domain should not be used as a standalone PPI prediction tool to identify virulence factors responsible for IAV infections [24]. Based on results of this work, IAV genomes across different strains comprise highly similar domains due to their similar structure (i.e., eight segments, encoding at least 11 proteins) and biological function. Furthermore, interactions that involve protein structures are facilitated not only by the protein domains, but also by various non-structured regions, such as interdomain linkers, N and C terminal structures or sequences, protein peptides [20]. Therefore, utilizing DISPOT exclusively may produce high number of false negative or false positive PPI predictions.

Mitochondria play an imperative role in antiviral innate immune response through the Mitochondiral antiviral-signaling protein (MAVS) *[MAVS_MOUSE (UniProt accession number: Q8VCF0)]* protein, a component of the Retinoic acid-inducible gene I (RIG-I) antiviral pathway. This pathway along with multiple others, is essential for combating and resolving viral infection, repair of damaged tissues, and generating adaptive immune response. It has been revealed that PB1-F2 inhibits antiviral cytokines and enhances expression of inflammatory cytokines through direct interaction with MAVS and other components of the RIG-I/MAVS system [18]. However, as protein sequences of both PB1-F2 and MAVS were not assigned any SCOP domain by SUPERFAMILY 2.0, it was not possible to verify this interaction via DISPOT.

A homeodomain-like domain was identified by DISPOT to be interacting with the viral protein domain present in the HA gene segment of IAV, with a statistical potential score of - 3.203 *(rounded to 3 d.p.).* However a study conducted by [12], which integrates both IAV-host PPIs detected using either small-scale or large-scale researches carried out experimentally or computationally found no evidence for an interaction with HA. In [12], homeobox protein MOX-2 (MEOX2) *[MEOX2_MOUSE, (UniProt accession number: P32443)]*, containing the homeodomain-like domain, was identified to be interacting with IAV gene segments PB1, PA, NA and M2 but not HA. By comparison, for DISPOT no interaction was detected between the homeodomain-like domain and segments PB1 and NA. Likewise, as protein sequences of segments PA and M2 were not assigned any SCOP domain by SUPERFAMILY 2.0, DISPOT could not be used to ascertain these interactions. Further, the cellular localisation for proteins with homeodomain-like domains was found to be in the nucleus according to UniProt, which indicates its interaction with HA would be unlikely, given that HA, mediates cell-surface recognition and viral entry.

Additionally, the reason for virulence levels to differ across mouse strains infected with the same IAV strain has not been uncovered as protein sequences of mouse strains retrieved from UniProt were derived from referencing the C57BL/6J mouse strain only. This limitation is because currently whole proteome sequences of other mouse strains (i.e., BALB/c, DBA/2 and FVB/J) are not available in any public database. Translation of strain specific genomic sequences to whole proteomes is a challenging task needing extensive experimental effort. In this work, obtaining these proteomes by means of experimental protein sequencing was not possible as the necessary materials and labor were not available.

### 5 FUTURE WORK

To bridge the gaps in this work, sequence-based PPI prediction methods can be employed to substantiate DDIs identified by DISPOT. An example is the Human-Virus Protein-Protein Interactions (HVPPI) web server, developed by X.Yang and colleagues [40]. HVPPI applied an unsupervised sequence embedding technique, doc2vec, to represent protein sequences as low-dimensional rich feature vectors. Then, a random forest classifier was trained using a training dataset that covers known PPIs between human and all viruses to predict human-virus PPIs. Lastly, the HVPPI web server automatically calculates the interaction probability of a query protein pair. The data to be used as input to HVPPI can be constructed as follows: Firstly, human protein sequences can be obtained using mouse and human homologs. Next, all protein sequences with SCOP domain(s) assigned to them can be trimmed, following the collected start and end residue numbers. Subsequently, trimmed protein sequences can be paired corresponding to DDIs recognized by DISPOT. Interaction probabilities provided as predicted outputs by HVPPI for each IAV-human protein pair can then be used to detect false positives. For protein sequences not assigned to any SCOP domain, complete protein sequences can be used, which will in turn aid with the detection of false negatives.

As an extension of this work from the raw dataset, instead of ignoring non-standard strains, the protein sequences of recombinant or mutant IAV strains can be reproduced via manually changing the protein sequences of wild-type or laboratory IAV strains that are available in UniProt. This enriches the dataset further.

The DISPOT statistical potentials, HVPPI interaction probabilities and LD_50_ values can be incorporated to represent the PPI network as a weighted undirected graph. Later, graph embedding methods can be applied to this weighted graph to learn low-dimensional node representations [42]. Structural information of PPI, such as the degree, position and neighbouring nodes in a graph has been recognized to be helpful in PPI prediction [39].

### 6 CONCLUSION

As IAV is a significant danger to global human health and life, it is critical to have deeper, accurate as well as reliable insights and knowledge on the virulence factors responsible for IAV infections to counteract potential outbreaks [15]. This work built upon a previously curated dataset of lethal dose studies of IAV infection in mice. Thereafter, superfamily domains involved in DDIs between IAV and mice were discovered, and ranked according to statistical interaction potentials calculated by DISPOT. A one-stop web server integrating information collated from literature and various databases, namely, NCBI Taxonomy, UniProt and SUPERFAMILY 2.0 with the DDI network was constructed. Furthermore, the web server is scalable and can seamlessly accommodate addition of new functions and data when future research is carried out.

## AUTHOR CONTRIBUTIONS

Ng and Rashid conceived and designed the method. Ng implemented the method. Ng and Rashid wrote the manuscript. Kwoh supervised throughout the conception, design, implementation and writing stages. All authors reviewed and approved the final manuscript.

## ACKNOWLEDGMENTS

The authors thank F.X. Ivan for constructing the initial dataset of infection records that was used to develop the virulence network here. They also thank the following consortia of databases and programs: NCBI PubMed and Taxonomy, UniProt, SUPERFAMILY 2.0 and DISPOT.

## SUPPLEMENTAL DATA

Supplementary data are available at Frontiers.

## DATA AVAILABILITY STATEMENT

The data and web server presented in this work are available at https://github.com/tengann/IAV-Host-PPI-Database and https://iav-ppi.herokuapp.com/home respectively.

