## Supplementary_Material for "Virulence Network of Interacting Influenza-Host Protein Domains"

### SUPPLEMENTARY FIGURES

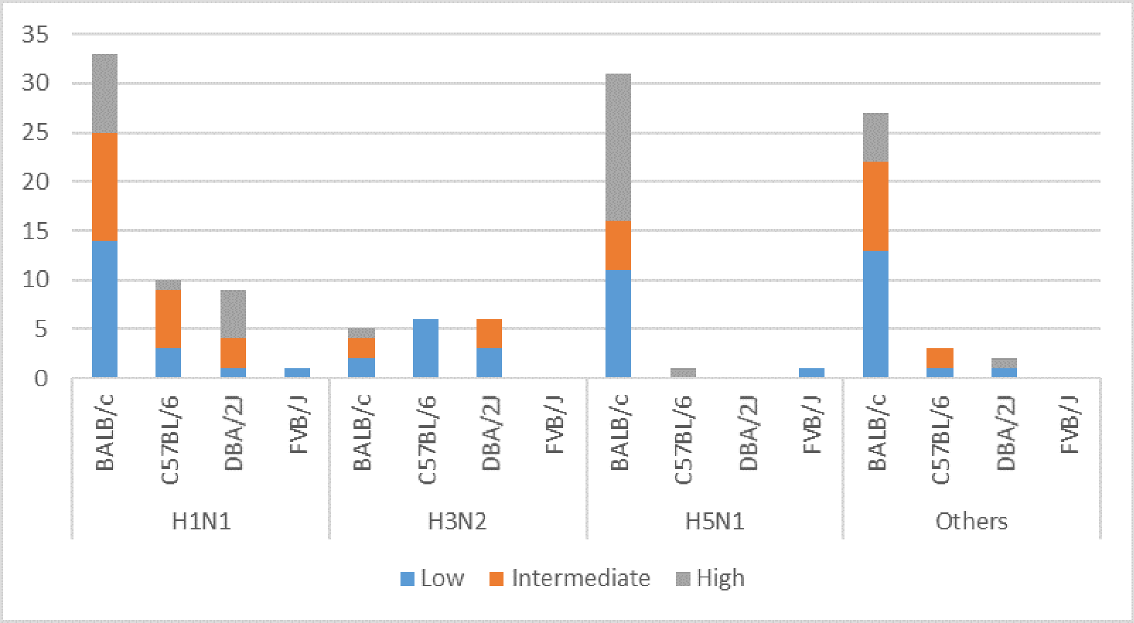


**Figure S1.** Cross-tabulation between IAV subtypes and mouse strains (based on cleaned dataset of 135 infection records), with breakdown according to three-class virulence classification problem. Number of cases classified as ‘virulent’ under the two-class problem is the sum of the number of ‘intermediate’ and ‘high’ in the three-class problem, while the number of cases classified as ‘avirulent’ is equivalent to the number of ‘low’ virulence cases. ‘Others’ refers to the aggregation of infection records from IAV subtypes - H1N2, H3N8, H5N2, H5N5, H5N6, H5N8, H7N2, H7N3, H7N7, H7N9.

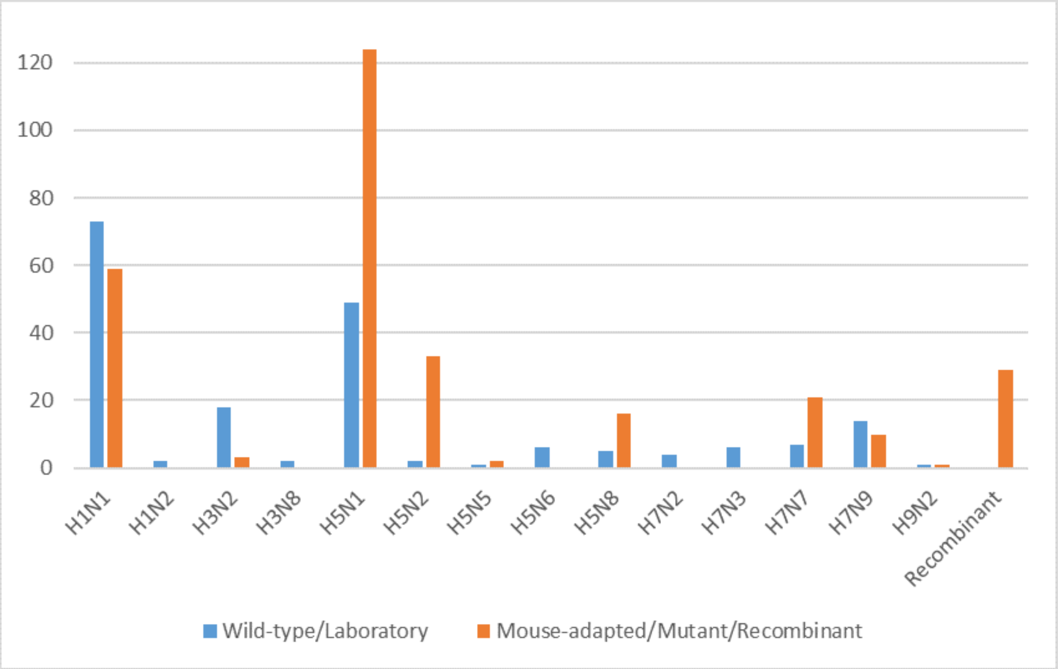


**Figure S2.** Initial dataset of 488 infection records - Proportion of wild-type/laboratory against mouse-adapted/mutant IAV strains. 'Recombinant' refers to IAV formed by the combination of protein segments retrieved from at least two different IAV subtypes. Infection records involving mouse-adapted, mutant and recombinant IAV strains (as represented by the orange bars) were first omitted, reducing the number of infection records to 190.

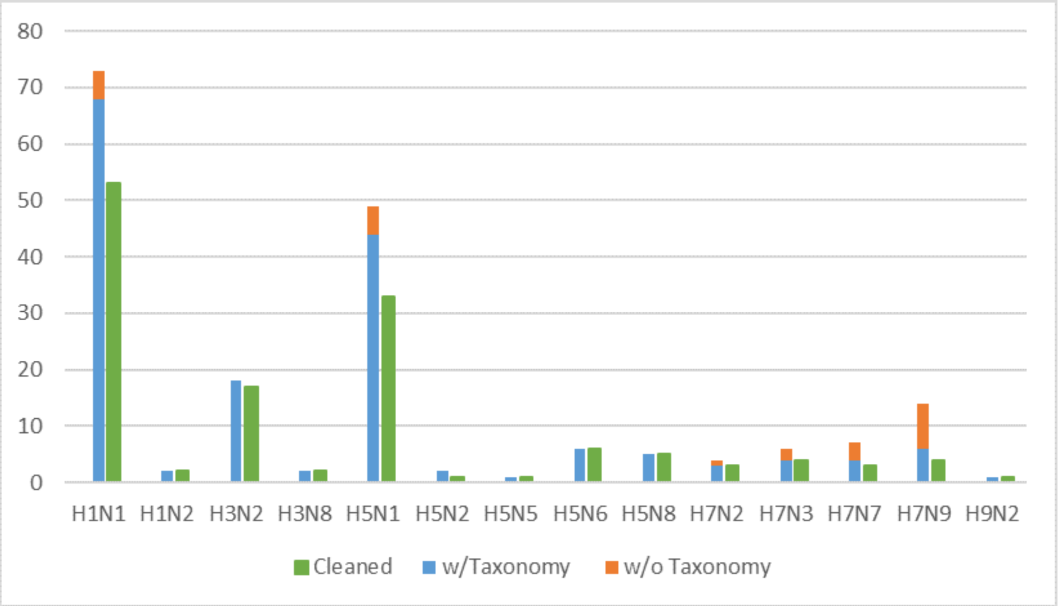


**Figure S3.** Proportion of all wild-type/laboratory IAV strains (separated into with and without Taxonomy ID) against remaining records in cleaned dataset. Infection records involving wild-type IAV where Taxonomy ID could not be found (orange bars) were then omitted, reducing the number of infection records to 166 (blue bars). 'Cleaned' refers to the final dataset of 135 infection records (green bars) available in IAV-Host PPI database, where each record relates to a unique combination of IAV and mouse strain.

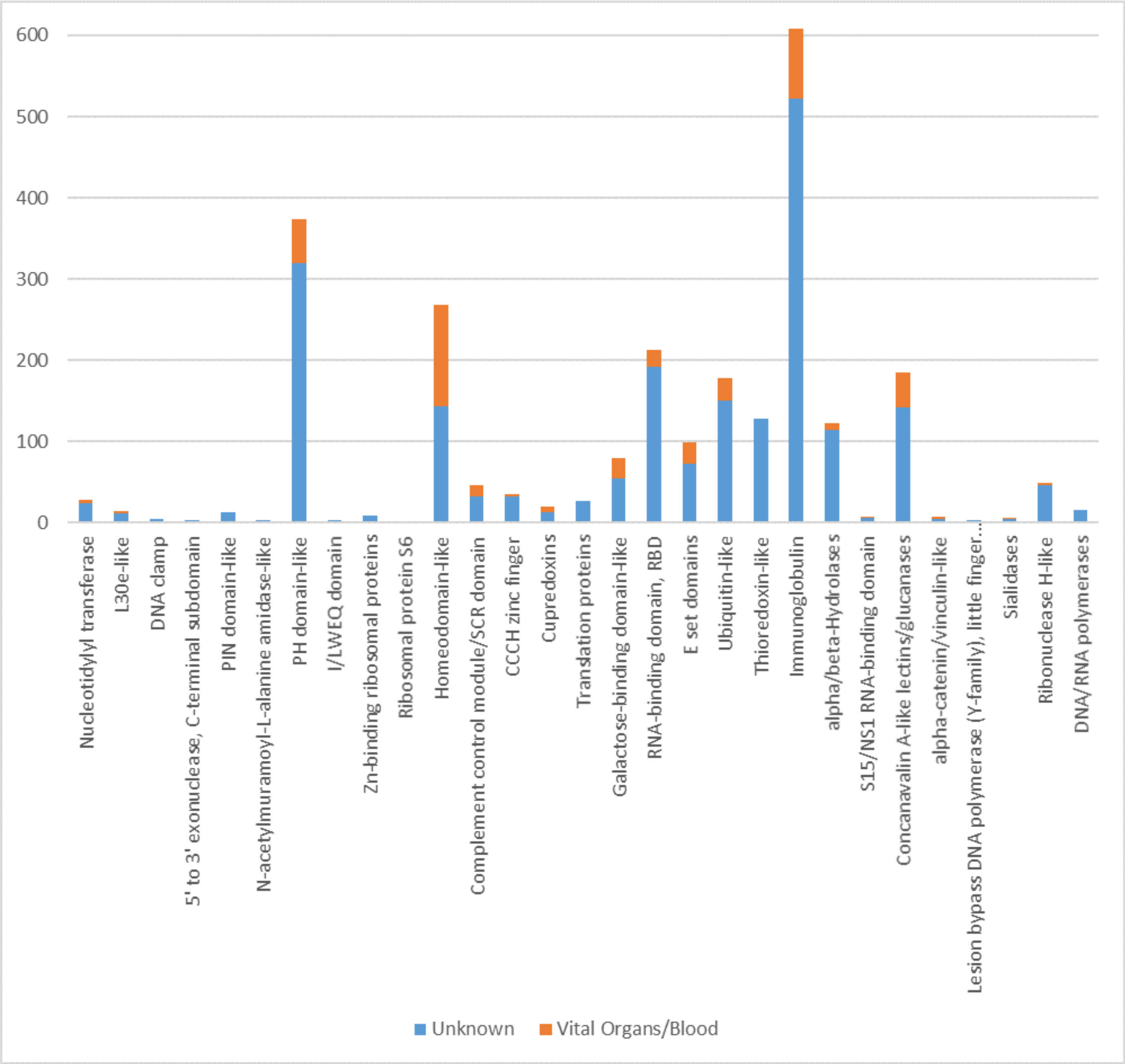


**Figure S4.** Mouse protein tabulation, grouped according to the 29 interacting SCOP superfamily. ‘Vital Organs/Blood’ refers to the aggregation of mouse proteins that can be found in a mouse’s lungs, brain, liver, kidney, spleen, heart and blood. ‘Unknown’ refers to the remaining mouse proteins, found in all other parts of a mouse’s body.
